## Supplementary materials for "Modulation of alpha oscillations by attention is predicted by hemispheric asymmetry of subcortical regions"

Table 1. Akaike Information Criterion (AIC) and Bayesian Information Criterion (BIC) values for all possible combinations of regressors (Lateralized Volume of subcortical structures). The selected model, with lowest AIC, is marked in green.

| *Regression* | **Comb. 1** | **Comb. 2** | **Comb. 3** | **Comb. 4** | **Comb. 5** | **Comb. 6** | **Comb. 7** | **Comb. 8** | **Comb. 9** | **Comb. 10** | **Comb. 11** | **Comb. 12** | **Comb. 13** | **Comb. 14** | **Comb. 15** | **Comb. 16** | **Comb. 17** | **Comb. 18** | **Comb. 19** | **Comb. 20** | **Comb. 21** | **Model criterion** |
|---|---|---|---|---|---|---|---|---|---|---|---|---|---|---|---|---|---|---|---|---|---|---|
| $HLM \sim \beta_0 + \beta_{LV_1}$ | -81.63 | -77.78 | -73.75 | -78.62 | -71.94 | -73.18 | -71.48 | NaN | NaN | NaN | NaN | NaN | NaN | NaN | NaN | NaN | NaN | NaN | NaN | NaN | NaN | **AIC** |
| | -77.33 | -73.48 | -69.45 | -74.32 | -67.64 | -68.88 | -67.18 | NaN | NaN | NaN | NaN | NaN | NaN | NaN | NaN | NaN | NaN | NaN | NaN | NaN | NaN | **BIC** |
| $HLM \sim \beta_0 + \beta_{LV_1} + \beta_{LV_2}$ | -86.72 | -79.93 | -82.66 | -77.88 | -78.04 | -78.00 | -76.42 | -81.27 | -75.82 | -75.83 | -73.89 | -77.41 | -70.41 | -71.36 | -69.94 | -74.79 | -76.05 | -74.63 | -70.08 | -68.23 | -69.30 | **AIC** |
| | -81.11 | -74.32 | -77.05 | -72.28 | -72.44 | -72.39 | -70.82 | -75.66 | -70.22 | -70.23 | -68.29 | -71.81 | -64.81 | -65.75 | -64.33 | -69.19 | -70.44 | -69.03 | -64.48 | -62.63 | -63.70 | **BIC** |
| | -85.30 | -87.54 | -82.79 | -82.96 | -82.64 | -81.00 | -76.18 | -76.19 | -76.29 | -78.86 | -79.00 | -78.73 | -74.25 | -74.24 | -74.32 | -79.67 | -74.61 | -74.06 | -72.55 | -78.51 | -78.88 | **AIC** |

| | | | | | | | | | | | | | | | | | | | | | | |
|---|---|---|---|---|---|---|---|---|---|---|---|---|---|---|---|---|---|---|---|---|---|---|
| $HLM \sim \beta_0 + \beta_{LV_1} + \beta_{LV_2} + \beta_{LV_3}$ | -78.46 | -80.71 | -75.95 | -75.13 | -75.81 | -74.17 | -69.34 | -69.35 | -69.46 | -72.02 | -72.17 | -71.90 | -67.42 | -57.40 | -67.49 | -72.84 | -67.77 | -67.22 | -65.72 | -71.68 | -72.05 | **BIC** |
| $HLM \sim \beta_0 + \beta_{LV_1} + \beta_{LV_2} + \beta_{LV_3} + \beta_{LV_4}$ | -85.52 | -81.42 | -81.27 | -81.21 | -83.50 | -83.73 | -83.30 | -79.33 | -78.80 | -78.85 | -77.26 | -77.72 | -77.09 | -72.43 | -72.54 | -72.49 | -75.15 | -74.95 | -75.04 | -70.57 | -76.78 | **AIC** |
| | -77.52 | -73.43 | -73.28 | -73.21 | -75.50 | -75.74 | -75.31 | -71.33 | -70.80 | -70.86 | -69.27 | -69.73 | -69.10 | -64.44 | -64.55 | -64.45 | -67.15 | -66.96 | -67.04 | -62.58 | -68.78 | **BIC** |
| $HLM \sim \beta_0 + \beta_{LV_1} + \beta_{LV_2} + \beta_{LV_3} + \beta_{LV_4} + \beta_{LV_5}$ | -81.51 | -81.90 | -81.30 | -77.65 | -77.41 | -77.17 | -79.95 | -79.32 | -79.49 | -75.29 | -73.88 | -73.36 | -73.75 | -68.77 | -71.22 | -75.31 | -72.78 | -73.87 | -68.85 | -72.63 | -67.74 | **AIC** |
| | -72.44 | -72.83 | -72.24 | -68.58 | -68.34 | -68.01 | -70.88 | -70.25 | -70.42 | -66.22 | -64.81 | -64.29 | -64.68 | -59.70 | -62.15 | -66.24 | -63.71 | -64.79 | -59.78 | -63.56 | -58.67 | **BIC** |
| $HLM \sim \beta_0 + \beta_{LV_1} + \beta_{LV_2} + \beta_{LV_3} + \beta_{LV_4} + \beta_{LV_5} + \beta_{LV_6}$ | -78.13 | -77.37 | -77.70 | -73.63 | -75.76 | -69.96 | -71.25 | NaN | NaN | NaN | NaN | NaN | NaN | NaN | NaN | NaN | NaN | NaN | NaN | NaN | NaN | **AIC** |
| | -68.07 | -67.30 | -67.63 | -63.56 | -65.70 | -59.89 | -61.19 | NaN | NaN | NaN | NaN | NaN | NaN | NaN | NaN | NaN | NaN | NaN | NaN | NaN | NaN | **BIC** |
| $HLM \sim \beta_0 + \beta_{LV_1} + \beta_{LV_2} + \beta_{LV_3} + \beta_{LV_4} + \beta_{LV_5} + \beta_{LV_6} + \beta_{LV_7}$ | -74.00 | NaN | NaN | NaN | NaN | NaN | NaN | NaN | NaN | NaN | NaN | NaN | NaN | NaN | NaN | NaN | NaN | NaN | NaN | NaN | NaN | **AIC** |
| | -63.01 | NaN | NaN | NaN | NaN | NaN | NaN | NaN | NaN | NaN | NaN | NaN | NaN | NaN | NaN | NaN | NaN | NaN | NaN | NaN | NaN | **BIC** |

| *Regression* | **Comb. 22** | **Comb. 23** | **Comb. 24** | **Comb. 25** | **Comb. 26** | **Comb. 27** | **Comb. 28** | **Comb. 29** | **Comb. 30** | **Comb. 31** | **Comb. 32** | **Comb. 33** | **Comb. 34** | **Comb. 35** | **Model criterion** |
|---|---|---|---|---|---|---|---|---|---|---|---|---|---|---|---|
| $HLM \sim \beta_0 + \beta_{LV_1}$ | NaN | NaN | NaN | NaN | NaN | NaN | NaN | NaN | NaN | NaN | NaN | NaN | NaN | NaN | **AIC** |
| | NaN | NaN | NaN | NaN | NaN | NaN | NaN | NaN | NaN | NaN | NaN | NaN | NaN | NaN | **BIC** |
| $HLM \sim \beta_0 + \beta_{LV_1} + \beta_{LV_2}$ | NaN | NaN | NaN | NaN | NaN | NaN | NaN | NaN | NaN | NaN | NaN | NaN | NaN | NaN | **AIC** |
| | NaN | NaN | NaN | NaN | NaN | NaN | NaN | NaN | NaN | NaN | NaN | NaN | NaN | NaN | **BIC** |

| | | | | | | | | | | | | | | | |
|---|---|---|---|---|---|---|---|---|---|---|---|---|---|---|---|
| $HLM \sim \beta_0 + \beta_{LV_1} + \beta_{LV_2} + \beta_{LV_3}$ | -77.34 | -74.59 | -71.90 | -71.93 | -73.54 | -75.43 | -73.45 | -68.27 | -66.72 | -67.51 | -72.40 | -70.89 | -72.05 | -66.31 | **AIC** |
| | -70.50 | -67.75 | -65.07 | -65.09 | -66.71 | -68.59 | -66.61 | -61.43 | -59.89 | -60.67 | -65.57 | -64.05 | -65.21 | -59.48 | **BIC** |
| $HLM \sim \beta_0 + \beta_{LV_1} + \beta_{LV_2} + \beta_{LV_3} + \beta_{LV_4}$ | -77.76 | -75.76 | -72.83 | -70.69 | -70.19 | -76.72 | -74.48 | -74.97 | -70.58 | -71.67 | -69.66 | -71.44 | -64.53 | -68.46 | **AIC** |
| | -69.76 | -67.77 | -64.84 | -62.70 | -62.19 | -68.73 | -66.49 | -66.98 | -62.59 | -63.68 | -61.67 | -63.44 | -56.53 | -60.47 | **BIC** |
| $HLM \sim \beta_0 + \beta_{LV_1} + \beta_{LV_2} + \beta_{LV_3} + \beta_{LV_4} + \beta_{LV_5}$ | NaN | NaN | NaN | NaN | NaN | NaN | NaN | NaN | NaN | NaN | NaN | NaN | NaN | NaN | **AIC** |
| | NaN | NaN | NaN | NaN | NaN | NaN | NaN | NaN | NaN | NaN | NaN | NaN | NaN | NaN | **BIC** |
| $HLM \sim \beta_0 + \beta_{LV_1} + \beta_{LV_2} + \beta_{LV_3} + \beta_{LV_4} + \beta_{LV_5} + \beta_{LV_6}$ | NaN | NaN | NaN | NaN | NaN | NaN | NaN | NaN | NaN | NaN | NaN | NaN | NaN | NaN | **AIC** |
| | NaN | NaN | NaN | NaN | NaN | NaN | NaN | NaN | NaN | NaN | NaN | NaN | NaN | NaN | **BIC** |
| $HLM \sim \beta_0 + \beta_{LV_1} + \beta_{LV_2} + \beta_{LV_3} + \beta_{LV_4} + \beta_{LV_5} + \beta_{LV_6} + \beta_{LV_7}$ | NaN | NaN | NaN | NaN | NaN | NaN | NaN | NaN | NaN | NaN | NaN | NaN | NaN | NaN | **AIC** |
| | NaN | NaN | NaN | NaN | NaN | NaN | NaN | NaN | NaN | NaN | NaN | NaN | NaN | NaN | **BIC** |

Table 2. The combination of structures for each regression model above. The selected model is marked in green.

| *Regression* | **Comb. 1** | **Comb. 2** | **Comb. 3** | **Comb. 4** | **Comb. 5** | **Comb. 6** | **Comb. 7** | **Comb. 8** | **Comb. 9** | **Comb. 10** | **Comb. 11** | **Comb. 12** | **Comb. 13** | **Comb. 14** | **Comb. 15** | **Comb. 16** | **Comb. 17** | **Comb. 18** | **Comb. 19** | **Comb. 20** | **Comb. 21** |
|---|---|---|---|---|---|---|---|---|---|---|---|---|---|---|---|---|---|---|---|---|---|
| $HLM \sim \beta_0 + \beta_{LV_1}$ | Th | CN | Put. | GP | Hipp. | Amyg. | Acc. | | | | | | | | | | | | | | |
| $HLM \sim \beta_0 + \beta_{LV_1} + \beta_{LV_2}$ | Th + CN | Th + Put. | Th + GP | Th + Hipp. | Th + Amyg. | Th + Acc. | CN + Put. | CN + GP | CN + Hipp. | CN + Amyg. | CN + Acc. | Put. + GP | Put. + Hipp. | Put. + Amyg. | Put. + Acc. | GP + Hipp. | GP + Amyg. | GP + Acc. | Hipp. + Amyg. | Hipp. + Acc. | Amyg. + Acc. |

| | | | | | | | | | | | | | | | | | | | | | |
|---|---|---|---|---|---|---|---|---|---|---|---|---|---|---|---|---|---|---|---|---|---|
| $HLM \sim \beta_0 + \beta_{LV_1} + \beta_{LV_2} + \beta_{LV_3}$ | Th + CN + Put. | Th + CN + GP | Th + CN + Hipp. | Th + CN + Amyg. | Th + CN + Acc. | Th + Put. + GP | Th + Put. + Hipp. | Th + Put. + Amyg. | Th + Put. + Acc. | Th + GP + Hipp. | Th + GP + Amyg. | Th + GP + Acc. | Th + Hipp. + Amyg. | Th + Hipp. + Acc. | Th + Amyg. + Acc. | CN + Put. + GP | CN + Put. + Hipp. | CN + Put. + Amyg. | CN + Put. + Acc. | CN + GP + Hipp. | CN + GP + Amyg. |
| $HLM \sim \beta_0 + \beta_{LV_1} + \beta_{LV_2} + \beta_{LV_3} + \beta_{LV_4}$ | Th + CN + Put. + GP | Th + CN + Put. + Hipp. | Th + CN + Put. + Amyg. | Th + CN + Put. + Acc. | Th + CN + GP + Hipp. | Th + CN + GP + Amyg. | Th + CN + GP + Acc. | Th + CN + Hipp. + Amyg. | Th + CN + Hipp. + Acc. | Th + CN + Amyg. + Acc. | Th + Put. + GP + Hipp. | Th + Put. + GP + Amyg. | Th + Put. + GP + Acc. | Th + Put. + Hipp. + Amyg. | Th + Put. + Hipp. + Acc. | Th + Put + Amyg. + Acc. | Th + GP + Hipp. + Amyg. | Th + GP + Hipp. + Acc. | Th + GP + Amyg. + Acc. | Th + Hipp. + Amyg. + Acc. | CN + Put. + GP + Hipp. |
| $HLM \sim \beta_0 + \beta_{LV_1} + \beta_{LV_2} + \beta_{LV_3} + \beta_{LV_4} + \beta_{LV_5}$ | Th + CN + Put. + GP + Hipp. | Th + CN + Put. + GP + Amyg. | Th + CN + Put. + GP + Acc. | Th + CN + Put. + Hipp. + Amyg. | Th + CN + Put. + Hipp. + Acc. | Th + CN + Put. + Amyg. + Acc. | Th + CN + GP + Hipp. + Amyg. | Th + CN + GP + Hipp. + Acc. | Th + CN + GP + Amyg. + Acc. | Th + CN + Hipp. + Amyg. + Acc. | Th + Put. + GP + Hipp. + Amyg. | Th + Put. + GP + Hipp. + Acc. | Th + Put. + GP + Amyg. + Acc. | Th + Put. + Hipp. + Amyg. + Acc. | Th + GP + Hipp. + Amyg. + Acc. | CN + Put. + GP + Hipp. + Amyg. | CN + Put. + GP + Hipp. + Acc. | CN + Put. + GP + Amyg. + Acc. | CN + Put. + Hipp. + Amyg. + Acc. | CN + GP + Hipp. + Amyg. + Acc. | Put. + GP + Hipp. + Amyg. + Acc. |
| $HLM \sim \beta_0 + \beta_{LV_1} + \beta_{LV_2} + \beta_{LV_3} + \beta_{LV_4} + \beta_{LV_5} + \beta_{LV_6}$ | Th + CN + Put. + GP + Hipp. + Amyg. | Th + CN + Put. + GP + Hipp. + Acc. | Th + CN + Put. + GP + Amyg. + Acc. | Th + CN + Put. + Hipp. + Amyg. + Acc. | Th + CN + GP + Hipp. + Amyg. + Acc. | Th + Put. + GP + Hipp. + Amyg. + Acc. | CN + Put. + GP + Hipp. + Amyg. + Acc. | | | | | | | | | | | | | | |
| $HLM \sim \beta_0 + \beta_{LV_1} + \beta_{LV_2} + \beta_{LV_3} + \beta_{LV_4} + \beta_{LV_5} + \beta_{LV_6} + \beta_{LV_7}$ | Th + CN + Put. + GP + Hipp. + Amyg. + Acc. | | | | | | | | | | | | | | | | | | | | |

| *Regression* | Comb. 22 | Comb. 23 | Comb. 24 | Comb. 25 | Comb. 26 | Comb. 27 | Comb. 28 | Comb. 29 | Comb. 30 | Comb. 31 | Comb. 32 | Comb. 33 | Comb. 34 | Comb. 35 |
|---|---|---|---|---|---|---|---|---|---|---|---|---|---|---|
| $HLM \sim \beta_0 + \beta_{LV_1}$ | NaN | NaN | NaN | NaN | NaN | NaN | NaN | NaN | NaN | NaN | NaN | NaN | NaN | NaN |
| $HLM \sim \beta_0 + \beta_{LV_1} + \beta_{LV_2}$ | NaN | NaN | NaN | NaN | NaN | NaN | NaN | NaN | NaN | NaN | NaN | NaN | NaN | NaN |
| $HLM \sim \beta_0 + \beta_{LV_1} + \beta_{LV_2} + \beta_{LV_3}$ | CN + GP + Acc. | CN + Hipp. + Amyg. | CN + Hipp. + Acc. | CN + Amyg. + Acc. | Put. + GP + Hipp. | Put. + GP + Amyg. | Put. + GP + Acc. | Put. + Hipp. + Amyg. | Put. + Hipp. + Acc. | Put. + Amyg. + Acc. | GP + Hipp. + Amyg. | GP + Hipp. + Acc. | GP + Amyg. + Acc. | Hipp. + Amyg. + Acc. |
| $HLM \sim \beta_0 + \beta_{LV_1} + \beta_{LV_2} + \beta_{LV_3} + \beta_{LV_4}$ | CN + Put. + GP + Hipp. | CN + Put. + GP + Amyg. | CN + Put. + GP + Acc. | CN + Put. + Hipp. + Amyg. | CN + Put. + Amyg. + Acc. | CN + GP + Hipp. + Amyg. | CN + GP + Hipp. + Acc. | CN + GP + Amyg. + Acc. | CN + Hipp. + Amyg. + Acc. | Put. + GP + Hipp. + Amyg | Put. + GP + Hipp. + Acc. | Put. + GP + Amyg. + Acc. | Put. + Hipp. + Amyg. + Acc | GP + Hipp. + Amyg. + Acc. |
| $HLM \sim \beta_0 + \beta_{LV_1} + \beta_{LV_2} + \beta_{LV_3} + \beta_{LV_4} + \beta_{LV_5}$ | NaN | NaN | NaN | NaN | NaN | NaN | NaN | NaN | NaN | NaN | NaN | NaN | NaN | NaN |
| $HLM \sim \beta_0 + \beta_{LV_1} + \beta_{LV_2} + \beta_{LV_3} + \beta_{LV_4} + \beta_{LV_5} + \beta_{LV_6}$ | NaN | NaN | NaN | NaN | NaN | NaN | NaN | NaN | NaN | NaN | NaN | NaN | NaN | NaN |

| | | | | | | | | | | | | | | |
|---|---|---|---|---|---|---|---|---|---|---|---|---|---|---|
| $HLM \sim \beta_0 + \beta_{LV_1} + \beta_{LV_2} + \beta_{LV_3} + \beta_{LV_4} + \beta_{LV_5} + \beta_{LV_6} + \beta_{LV_7}$ | NaN | NaN | NaN | NaN | NaN | NaN | NaN | NaN | NaN | NaN | NaN | NaN | NaN | NaN |

Figure 1. Lateralization volume of thalamus, caudate nucleus and globus pallidus in relation to hemispheric lateralization modulation of rapid invisible frequency tagging (HLM(RIFT)) on the right and behavioural asymmetry on the left. A and E, The beta coefficients for the best model (having three regressors) associated with a generalized linear model (GLM) where lateralization volume (LV) values were defined as explanatory variables for HLM(RIFT) (A) and behavioural asymmetry (E). Error bars indicate standard errors of mean (SEM). B and F, Partial regression plot showing the association between $LV_{Th}$ and HLM(RIFT) (B, p-value = 0.59) and behavioural asymmetry (F, p-value = 0.38) while controlling for $LV_{GP}$ and $LV_{CN}$. C and G, Partial regression plot showing the association between $LV_{GP}$ and HLM(RIFT) (C, p-value = 0.16) and behavioural asymmetry (G, p-value = 0.80) while controlling for $LV_{Th}$ and $LV_{CN}$ . D and H, Partial regression plot showing the association between $LV_{CN}$ and HLM(RIFT) (D, p-value = 0.53) and behavioural asymmetry (H, p-value = 0.74) while controlling for $LV_{Th}$ and $LV_{GP}$. Negative (or positive) LVs indices denote greater left (or right) volume for a given substructure; similarly negative HLM(RIFT) values indicate stronger modulation of RIFT power in the left compared with the right hemisphere, and vice versa; positive behavioural asymmetry value shows higher accuracy when the target was on the right as compared with left, and vice versa for negative behavioural asymmetry values. The dotted curves in B, C, D, F, G, and H indicate 95% confidence bounds for the regression line fitted on the plot in red.

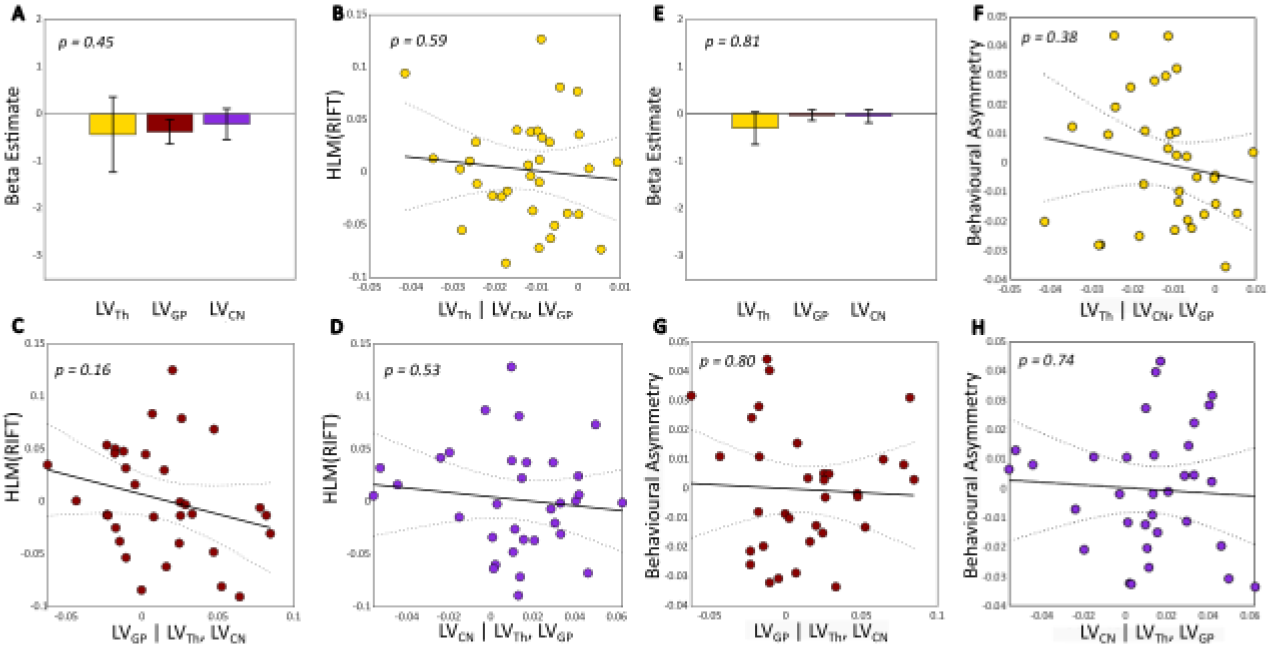


Table 3. Bayes factors for correlation between hemispheric laterality of subcortical structures with hemispheric lateralization modulation of rapid invisible frequency tagging (HLM(RIFT)) and with behavioural asymmetry (BA). The Pearson correlation between each subcortical structure with HLM(RIFT) and behavioural asymmetry was calculated. The likelihood of the data under the alternative hypothesis (the evidence of correlation) were subsequently compared to the likelihood under null hypothesis (absence of correlation), given the data. As it is demonstrated in the table, all Bayes factors were below or very close to 1 indicating evidence for the null hypothesis.

| Subcortical structure | HLM(RIFT) | BA |
|---|---|---|
| Thalamus | 0.14 | 0.20 |
| Caudate Nucleus | 0.18 | 0.15 |
| Putamen | 0.14 | 0.14 |
| Globus Pallidus | 0.34 | 0.13 |
| Hippocampus | 0.19 | 0.25 |
| Amygdala | 0.54 | 1.18 |
| Nucleus Accumbens | 0.16 | 0.17 |
